# Dimensionality reduction simplifies synaptic partner matching in an olfactory circuit

**DOI:** 10.1101/2024.08.27.609939

**Authors:** Cheng Lyu, Zhuoran Li, Chuanyun Xu, Kenneth Kin Lam Wong, David J. Luginbuhl, Colleen N. McLaughlin, Qijing Xie, Tongchao Li, Hongjie Li, Liqun Luo

## Abstract

The distribution of postsynaptic partners in three-dimensional (3D) space presents complex choices for a navigating axon. Here, we discovered a dimensionality reduction principle in establishing the 3D glomerular map in the fly antennal lobe. Olfactory receptor neuron (ORN) axons first contact partner projection neuron (PN) dendrites at the 2D spherical surface of the antennal lobe during development, regardless of whether the adult glomeruli are at the surface or interior of the antennal lobe. Along the antennal lobe surface, axons of each ORN type take a specific 1D arc-shaped trajectory that precisely intersects with their partner PN dendrites. Altering axon trajectories compromises synaptic partner matching. Thus, a 3D search problem is reduced to 1D, which simplifies synaptic partner matching and may generalize to the wiring process of more complex brains.

## Main Text

Proper function of the brain requires precise assembly of neural circuits during development. Advances in electron microscopy have been recently leveraged to dissect regional or even brain-wide connectivity patterns in an increasing number of species, from *C. elegans* to mammals (White et al., 1986; Briggman et al., 2011; Loomba et al., 2022; Schlegel et al., n.d.), revealing unprecedented precision of neural circuit wiring. Understanding how neural circuits establish such precise synaptic connections is a central goal of neurobiology.

A fundamental open question in synaptic partner selection is how to minimize the choice for a navigating axon. Axon guidance mitigates this problem by guiding an axon to a specific brain region (Dickson, 2002; Kolodkin & Tessier-Lavigne, 2011). After arriving at the appropriate brain region, how does an axon navigate the local 3D space to find its partners among many non-partners? Some neural circuits reduce this task load by organizing target selection in lower dimensions. For example, layered organization in vertebrate retina and fly optic lobe enables target selection to be restricted within a specific 2D layer after axons are targeted to a specific layer (Agi et al., 2024; Sanes & Zipursky, 2020). Here, even though strictly speaking the targets still occupy a 3D physical space and an axon moves in a 3D physical space, we can approximate these target selection problems as 2D since each axon only needs to search on a 2D plane to find its synaptic partner(s). But for many brain structures in which synaptic partners appear to be distributed in all three dimensions, an axon would need to code for the correct movement along each of the three axes as well as contain the capacity for recognizing all potential synaptic partners within that brain region, being attracted to the correct partners and repelled from the incorrect ones. Are there means to simplify the synaptic partner matching problem? Here, we illustrate that in the wiring of the olfactory circuit in the fly antennal lobe, the task complexity of synaptic partner searching is reduced from 3D to 1D.

### PN dendrites extend to the antennal lobe surface during development

In adult *Drosophila*, ∼50 types of olfactory receptor neurons (ORNs) synapse with 50 types of projection neurons (PN) in a one-to-one fashion at 50 discrete glomeruli. Each glomerulus forms a functional unit and the 50 glomeruli together occupy stereotyped 3D positions in the antennal lobe, with some exposed to the antennal lobe surface while others exclusively interior (Fig. 1, A and B). Previous studies show that the assembly of the fly olfactory circuit takes sequential steps. PN dendrites first elaborate and form a coarse map (Jefferis et al., 2004; Wong et al., 2023). ORN axons then circumnavigate ipsi- and contralateral antennal lobes from ∼18 hours to ∼32 hours after puparium formation (h APF) (Li et al., 2021; Okumura et al., 2016). Concomitant with ORN axons extending towards the contralateral antennal lobe, they initiate branches in the ipsilateral antennal lobe to search for their partner PN dendrites (Li et al., 2021; Xu et al., 2024) (Fig. 1A). However, the strategy an ORN axon employes to search for and match with partner PN dendrites remains incompletely understood. Does each ORN axon need to search the entire 3D space and scan through all the 50 PN types, or are there ways to reduce the number of PN candidates for each ORN type? Specifically, we note that in the vertebrate olfactory system, glomeruli are located on the surface of the olfactory bulb, simplifying target selection of ORN axons to a 2D problem (Mori & Sakano, 2011). Could a similar strategy be used in the developing *Drosophila* antennal lobe?

**Fig. 1.**
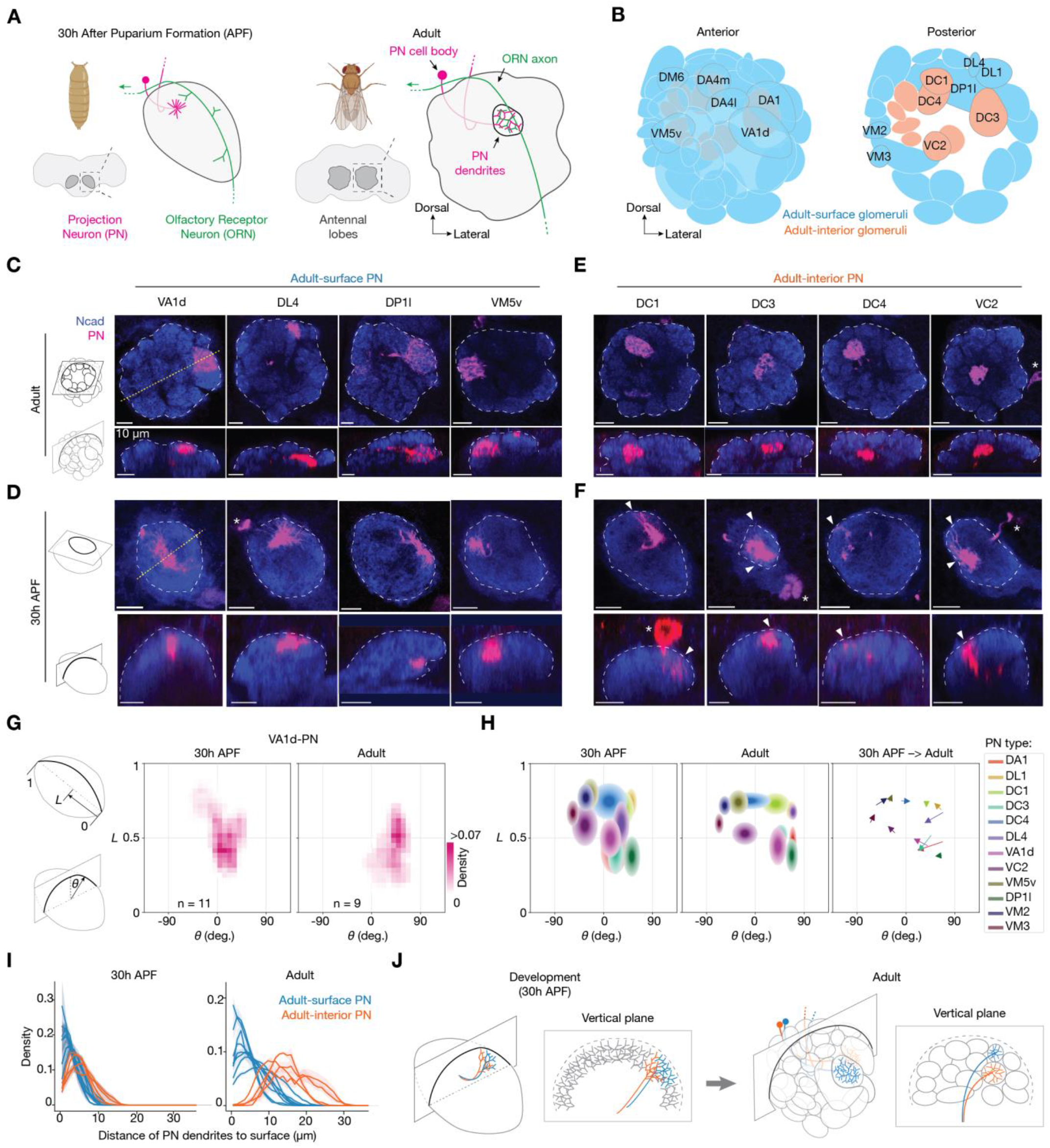
During development, PN dendrites are exposed to the antennal lobe surface regardless of their position in adults. (**A**) *Drosophila* brain and antennal lobe schematics, at 30 hours after puparium formation (h APF) (left) and adults (right). Antennal lobes are highlighted in dark grey surrounded by dash squares and magnified to the right. At 30h APF, PN dendrites (magenta) innervate similar positions as in adults and ORN axons (green) navigate along the surface of the ipsilateral antennal lobe from entry point at the bottom right towards midline at the top left. In adults, ORNs and PNs establish one-to-one connections in individual glomeruli that form a 3D glomerular map. (**B**) Adult antennal lobe schematic with ∼50 glomeruli circled. Cyan: glomeruli located at the surface of the antennal lobe; orange: glomeruli located in the interior of the antennal lobe. (**C**) Optical sections showing dendrites of specific adult-surface PN types (magenta, labeled by a membrane-targeted GFP driven by separate genetic drivers specific to PN types labeled above) viewed from the horizontal plane (top row) and the vertical plane (bottom row) of the antennal lobe in adults. White dash lines outline the antennal lobe neuropil stained by the N-cadherin (NCad) antibody (blue). Yellow dotted lines indicate the intersections with the vertical planes shown below. Vertical planes are reconstructed from 3D image volumes where optical sections were taken horizontally. The top and bottom rows show the same brains. Scale bar = 10 *µ*m in this and all other panels. (**D**) Same as (C), but with data from 30h APF. (**E** and **F**) Same as (C) and (D), but for adult-interior PN types. In (F), arrowheads indicate PN dendrites extending to the antennal lobe surface. *, PN cell body. (**G**) Probability distribution of VA1d-PN dendritic pixels projected onto the antennal lobe surface during development (middle) and in adults (right). The 2D antennal lobe surface is flattened and decomposed into two axes: the *x*-axis indicates the angle *θ* of each vertical plane and the *y*-axis indicates the position *L* along the long axis of the antennal lobe. Schematic definition of *θ* and *L* on the left. (**H**) Probability distribution of dendritic pixels from twelve PN types projected onto the antennal lobe surface. Left and middle, each ellipse corresponds to one PN type, with ellipse centers matching PN-dendrite centroids, and ellipse boundaries matching the standard deviations of PN dendrites along the *x*- and *y*-axes, respectively. Right, arrows represent the shift of centroid of the same-type ellipses from 30h APF to adults. See Fig. S1 for the n of each group. (**I**) Probability distribution of the shortest distance in 3D space from PN dendritic pixels to the antennal lobe surface during development (left) and in adults (right). Each line represents data from an individual PN type, population mean ± s.e.m. (**J**) Schematics of two individual PN types during development (left) and in adults (right), viewed from +45° anterior and from a single vertical plane. Note that PN dendrites extend to the antennal lobe surface during development regardless of their surface-or-interior positions in adults.

To address these questions, we began by examining the distribution of PN dendrites during development with single-type resolutions. We generated a collection of genetic drivers that label single PN types across developmental stages (Fig. 1, C–F and Fig. S1), using split-GAL4 (Luan et al., 2006) and Flp/FRT-based intersection strategies (Golic & Lindquist, 1989). Using these genetic drivers, we compared the dendritic patterns of single PN types between the adult and the developmental stages when ORN axons start navigating the antennal lobe. For PNs that innervate surface glomeruli in the adult antennal lobe (referred to as adult-surface PNs below), during development, their dendrites extended to the antennal lobe surface (Fig. 1, C and D). Surprisingly, for PNs innervating the interior glomeruli in the adult antennal lobe (referred to as adult-interior PNs below), during development, some of their dendrites also extended to the antennal lobe surface (Fig. 1, E and F).

Quantitative analyses revealed that the dendritic locations of all PN types at 30h APF approximated their future glomerular positions in adults (Fig. 1, G and H, Fig. S1). Furthermore, dendritic distributions along the radius of the antennal lobe confirmed that all four adult-interior PN types we tested extended a portion of their dendrites to the antennal lobe surface, just like the eight adult-surface PN types (Fig. 1I). The surface extension of adult-interior PN dendrites during development is unlikely a result of cognate ORN and PN interactions, as the surface extension was also observed in the same PN types at an earlier developmental stage before ORN axons had reached the antennal lobe (Fig. S2). Thus, both adult-surface and adult-interior PNs extend their dendrites to the antennal lobe surface during development (Fig. 1J). This raises a hypothesis that ORNs and PNs establish synaptic partners on the 2D antennal lobe surface during development, instead of the 3D antennal lobe volume seen in adults. To test this hypothesis, we next examined ORN axons during development.

### ORN axons take cell-type specific trajectories at the antennal lobe surface during development

To facilitate the encountering of correct partners, each ORN axon should ideally navigate along a trajectory that intersects with the dendrites of cognate PNs. To test this, we generated a collection of genetic drivers that label single-type or groups of ORNs across developmental stages (Fig. 2, A and B; Figs. S3 and S4), using approaches similar to the generation of specific PN drivers. When viewing from a vertical perspective orthogonal to navigating axons, we found that all ORN axons navigated along the spherical surface of the antennal lobe, regardless of their surface-or-interior positions in adults (Fig. 2B and Fig. S4). That ORN axons navigate along the surface of the antennal lobe is consistent with previous studies examining axons in bulk (Joo et al., 2013; Okumura et al., 2016).

**Fig. 2.**
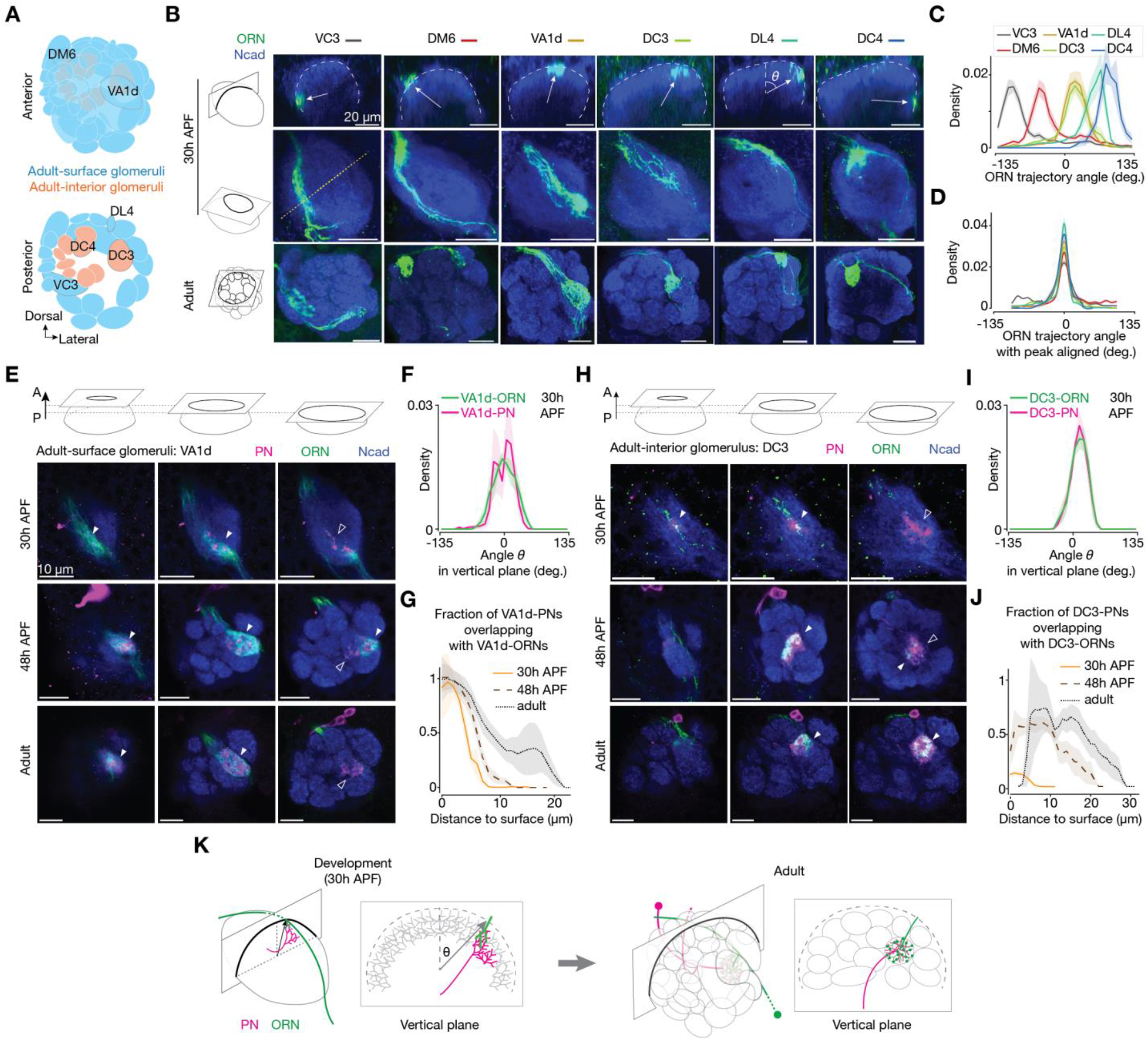
During development, ORN axons take cell-type specific trajectories and contact cognate PN dendrites first on the antennal lobe surface. (**A**) Adult antennal lobe schematic highlighting six glomeruli, corresponding to the six ORN types shown in (B– H). (**B**) Single ORN types (green, labeled by a membrane-targeted GFP driven by separate genetic drivers) at 30h APF (top and middle rows, same brains) and in adults (bottom rows). Top, single optical section from vertical plane with dash lines outlining the antennal lobe neuropil stained by the N-cadherin antibody (blue). Reconstructed from 3D image volumes where optical sections were taken horizontally. Arrows point from the antennal lobe center to the average positions of ORN axons. Trajectory angle *θ* is defined in the DL4 panel. Middle and bottom, maximum projection of horizontal optical sections of antennal lobes at 30h APF and adults, respectively. The yellow dotted line indicates the intersection with the vertical plane shown above. Scale bar = 20 *µ*m. (**C**) Probability distribution of the axon’s angular position from single-type ORNs. Population mean ± s.e.m. For all genotypes, n ≥ 9. (**D**) Same as (C), but with each data curve aligned to its peak to minimize data variance between brains and more accurately reflect the width of the probability distribution. (**E**) Single optical section showing VA1d-ORNs (green, labeled by membrane-targeted GFP driven by a split-GAL4) and VA1d-PNs (magenta, labeled by membrane-targeted RFP driven by a split-LexA (Ting et al., 2011)). From left to right, anterior, middle, and posterior sections from the same brain. From top to bottom, data from 30h APF, 48h APF, and adults. Filled arrowheads indicate examples where ORN axons and PN dendrites overlap. Open arrowheads indicate examples where PN dendrites do not overlap with ORN axons. Scale bar = 10 µm in this and all other panels. (**F**) Probability distribution of the angular position of VA1d-ORNs and VA1d-PNs. Same definition of the angle *θ* as in (C). Only vertical planes with PN dendrites were analyzed. Population mean ± s.e.m.; n = 9. (**G**) Fraction of VA1d-PNs overlapping with VA1d-ORNs, as a function of the distance from PN pixels to antennal lobe surface. For a given distance on the *x*-axis, *y* value of 1 means that all the VA1d-PN dendrites within that distance bin match with VA1d-ORN axons. Population mean ± s.e.m. For all time points, n ≥ 8. (**H–J**) Same as (E–G), but with data from DC3-ORNs and DC3-PNs. For all groups, n = 12. (**K**) Schematics of the same ORN-PN pair during development (left) and in adults (right). Note that DC3-ORN axons and DC3-PN dendrites are present at the antennal lobe surface during development but not in adults.

Interestingly, axons of each of the 6 ORN types we examined took a specific angular trajectory when viewed from a vertical perspective (Fig. 2, B–D), consistent with the positioning of their axonal tracts in adults. Some ORN types could have substantial trajectory overlaps, as observed in DC3-ORNs and VA1d-ORNs.

Axons from complementary ORN groups together covered the entire anterior surface of the antennal lobe (Fig. S4). These findings echo with PN dendrites extending to specific locations at the antennal lobe surface during development, and suggests that partner PN dendrites and ORN axons first meet on the 2D antennal lobe surface.

### ORN axons first contact cognate PN dendrites at the antennal lobe surface

To test whether the precise locations of PN dendrites and ORN axons enable future synaptic partners to be closer to each other, we labeled individual ORN types and their cognate PN types with different markers in the same brain across developmental stages (Fig. 2, E and H). Consistently, we observed that PN dendrites occupied a similar narrow angular range, coinciding precisely with the axons of cognate ORNs (Fig. 2, F and I). For an adult-surface glomerulus, VA1d, ORNs first contacted PNs at the antennal lobe surface (Fig. 2E top row, Fig. 2G orange curve) and maintained this trend to adults (Fig. 2E bottom rows, Fig. 2G black curves).

Similarly, for an adult-interior glomerulus, DC3, even though in adults the matching ORN axons and PN dendrites were in interior antennal lobe (Fig. 2H bottom rows, Fig. 2J black curves), ORN axons also first contacted PN dendrites at the surface during development (Fig. 2H top row, Fig. 2J orange curve).

Altogether, these results suggest a working model where individual ORN types and their cognate PNs first contact at the antennal lobe surface, regardless of their surface-or-interior positions in adults (Fig. 2K). We previously found that contact between cognate ORN and PN branches during development correlates with higher activity of filamentous actin, leading to stabilization of transient ORN axon branches (Xu et al., 2024). This suggests the overlaps we observed between ORN axon branches and PN dendrites at the surface participate in synaptic partner selection. Below, we further tested this using genetic perturbation experiments.

### Adult-interior PNs leave more dendrites on the surface after missing cognate ORNs

If the surface-located branches from adult-interior PNs are indeed expecting ORN partners during development, then missing the ORN partners during this time window may cause these PN branches to stay at the surface, perhaps connecting with other ORN types (Fig. 3, A and B). We tested this hypothesis by genetically altering ORN trajectories in two different adult-interior glomeruli where genetic reagents were available (Fig. 3, C–K). Our previous studies show that Sema-2b (Joo et al., 2013) and Toll-family proteins (Ward et al., 2015) are differentially expressed in ORN axons along the medial-lateral axis orthogonal to the trajectories ORN axons take to navigate across the antennal lobe surface, and that Sema-2b instructs trajectory choice of ORN axons (Joo et al., 2013). Using genetic drivers that label different ORN groups, we confirmed that manipulating the expression level of these proteins could alter the ORN trajectory during development in a way consistent with their expression patterns in the antennal lobe (Fig. S5). Therefore, in all genetic perturbation experiments below, we rerouted specific ORN axons by combinatorially manipulating the expression levels of Sema-2b and Toll proteins in specific ORN types. In wild-type adults, DC4-ORN axons match DC4-PN dendrites near the center of the antennal lobe (Fig. 3D). After we experimentally rerouted most DC4-ORN axons during development, DC4-PNs no longer matched DC4-ORNs and a portion of their dendrites remained at the antennal lobe surface in adults (Fig. 3E and Fig. S6; see Table S1 for detailed genotypes). Quantitative analysis revealed that surface dendrites in adults remained non-zero, unlike controls (Fig. 3F). We made similar findings for another adult-interior glomerulus, DC3 (Fig. 3, G, H and K).

**Fig. 3.**
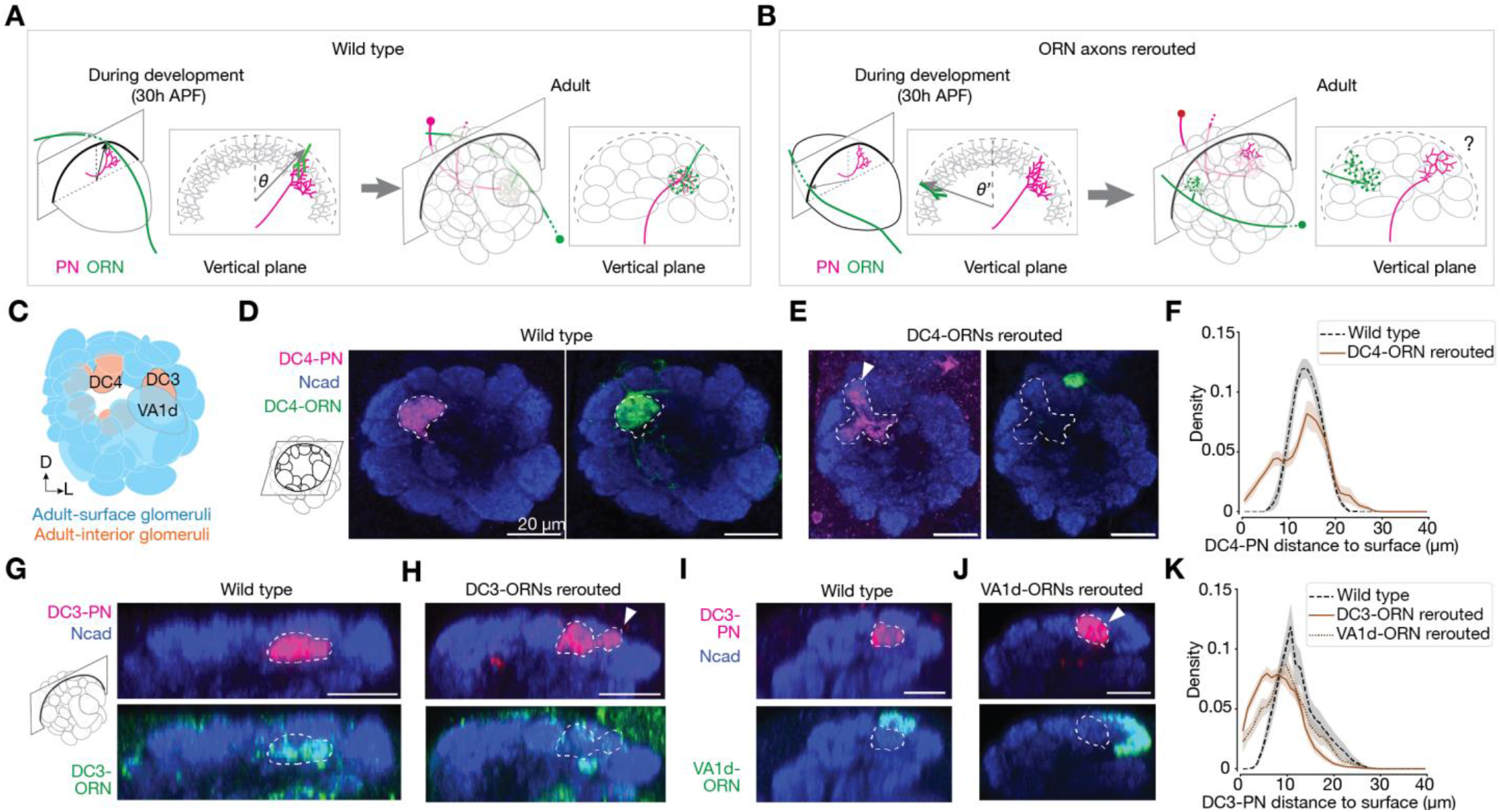
Dendrites of adult-interior PNs remain at the antennal lobe surface in adults when most axons of cognate ORNs are rerouted during development. (**A**) Schematics of the same ORN-PN pair during development (left) and in adult (right). Same as Fig. 2K. (**B**) Same as (A), but with ORN axons largely rerouted and missing cognate PNs during development (left). This could lead to adult-interior PNs remain at the surface in adult (right, indicated by a question mark). (**C**) Adult antennal lobe schematic labeling three glomeruli, corresponding to the three ORN-PN pairs shown in (D–J). Some glomeruli were omitted for visualization clarity. (**D**) Single optical section of DC4-ORNs (green, labeled by membrane-targeted GFP driven by a split-GAL4) and DC4-PNs (magenta, labeled by membrane-targeted RFP driven by a split-LexA) in a wild-type brain. Dashed lines outline the boundary of PN dendrites. Scale bar = 20 *µ*m in this and all other panels. (**E**) Same as (D), but with the trajectory of DC4-ORN axons changed through genetic manipulations (Toll-7 overexpression; see Fig. S6). The arrowhead indicates DC4-PNs innervating the antennal lobe surface. (**F**) Probability distribution of the distance from DC4-PN dendritic pixels to the antennal lobe surface in 3D space. Mean ± s.e.m. For all genotypes, n ≥ 6. (**G** and **H**) Same as (D) and (E), but with DC3-ORNs and DC3-PNs shown in a vertical plane and a different genetic manipulation (*Toll-6* and *Toll-7* RNAi; see Fig. S6). (**I** and **J**) Same as (G) and (H), but with the trajectory of VA1d ORNs changed through genetic manipulations (*Sema-2b* RNAi and *Toll-7* RNAi; see Fig. S6). (**K**) Same as (F), but with data from DC3-PNs. For all genotypes, n ≥ 11.

These results suggest that early in development, adult-interior ORN axons and PN dendrites make contact and form connections at the antennal lobe surface and together move towards the antennal lobe center as development proceeds. To examine the nature of the force that drives this inward movement, we genetically rerouted axons of VA1d-ORNs—adult-surface ORNs innervating the VA1d-glomerulus exterior to the DC3-glomerulus (Fig. 3, C and I, Fig. S6). This rerouting caused DC3-PN dendrites to remain at the antennal lobe surface in adults, unlike controls (Fig. 3, J and K, Fig. S6). This suggests that adult-interior ORNs and PNs first establish connections at the surface and their neurites are pushed towards the center by neurites from nearby adult-surface glomeruli.

### The accuracy of ORN-PN matching correlates with the accuracy of ORN trajectories

Summarizing the results so far, we found that, during development, the synaptic partner selection process of a 3D antennal lobe (Fig. 4A) occurs on a 2D antennal lobe surface. Dendrites of individual PN types either reside at (adult-surface types) or extend to (adult-interior types) specific parts of the antennal lobe surface, and axons of individual ORN types sweep through the antennal lobe surface, taking specific trajectories that precisely intersect with the dendrites of their partner PNs (Fig. 4B; see Fig. 2, F and I). This way, growing ORN axons can simply scan a thin stripe of the antennal lobe surface to identify PN partners, instead of the entire volume. A working model emerges that further reduces the dimensionality of partner selection for each ORN type from 2D to 1D: each ORN type only searches for synaptic partners within a 1D narrow stripe nearby its axon trajectory and all the ORN types altogether cover the entire antennal lobe surface (Fig. 4C).

**Fig. 4.**
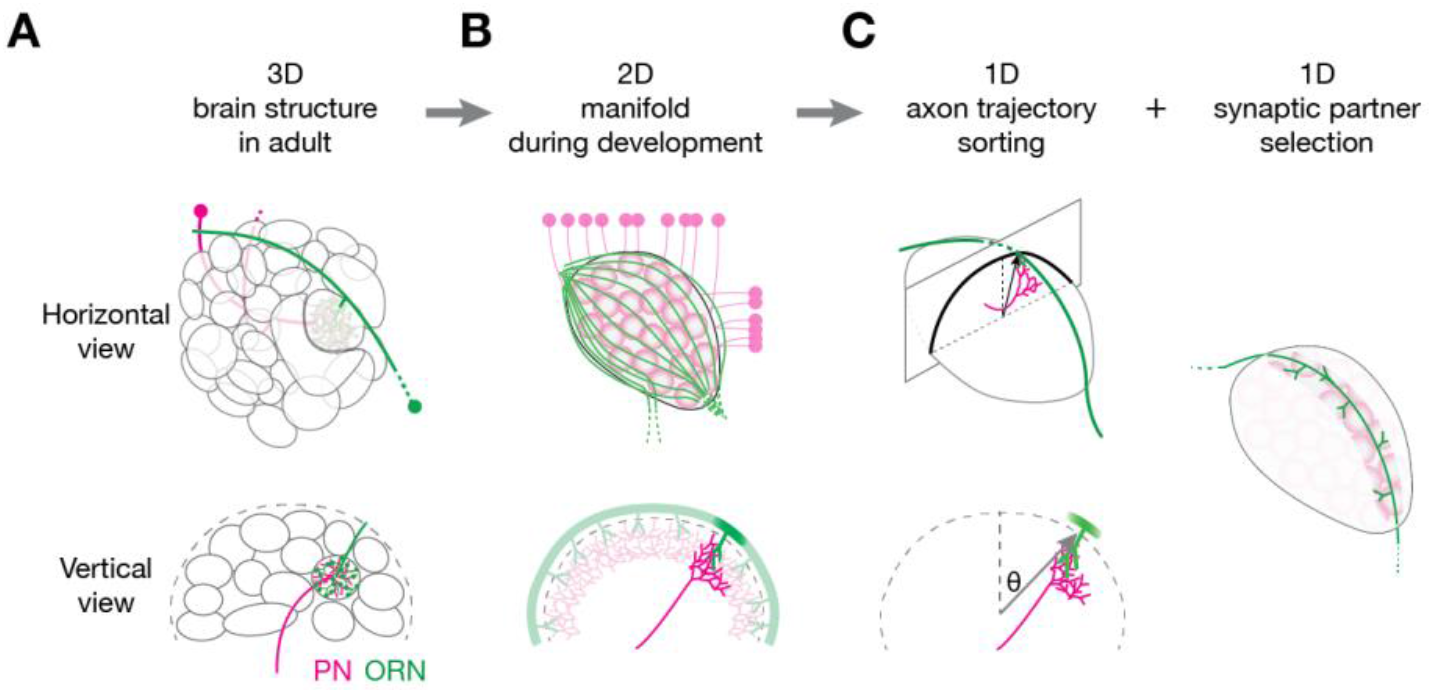
Summary of dimensionality reduction in the ORN-PN synaptic partner matching process. (**A**) The distribution of glomeruli appears 3D in adults. (**B**) During development, both PN dendrites and ORN axons search for partners at the 2D antennal lobe surface. (**C**) The search space for an individual ORN type is further reduced to 1D because their axons follow a specific trajectory on the 2D antennal lobe surface. See text for detail.

One way to further test this working model is to examine how the accuracy of ORN axon trajectory affects the accuracy of ORN-PN partner matching. If each ORN type only searches within the vicinity of its trajectory, then changing its trajectory should impair ORN-PN partner matching, with the degree of trajectory deviation determining the degree of mismatching. Using genetic drivers that label specific ORN types across developmental stages, we performed genetic manipulations in specific ORN types and altered their trajectories to different degrees in both directions during development. We then examine the accuracy of ORN-PN partner matching by labeling cognate ORNs and PNs with distinct markers in the same adult brain (Fig. 5).

**Fig. 5.**
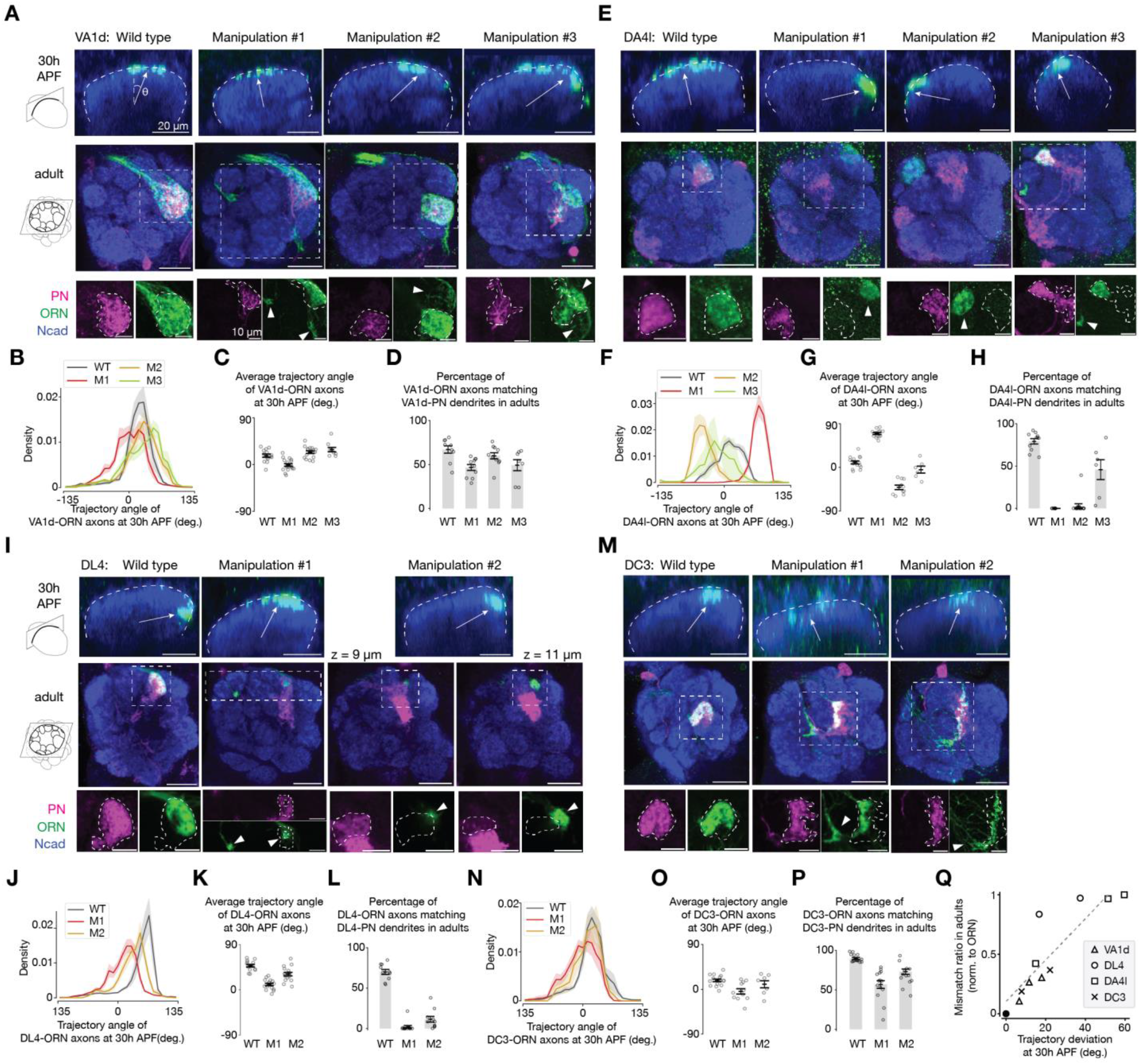
The accuracy of ORN-PN synaptic partner matching correlates with the accuracy of ORN trajectories. (**A**) Single optical sections showing VA1d-ORN axons from a vertical view during development (top) and horizontal view in adult (middle and bottom). Top, dash lines outline the antennal lobe neuropil. Arrows point from the antennal lobe center to the average positions of ORN axons. Images in the bottom row is a zoom-in from the dashed squares in the middle row. Bottom, dashed lines outline the boundary of PN dendrites. Arrowheads indicate ORN axons mismatching with cognate PN dendrites. Left column represents wild-type condition, other columns represent the three manipulation conditions: (1) *Sema-2b* RNAi; (2) *Toll-7* RNAi; (3) *Sema-2b* RNAi and *Toll-7* RNAi. See Table S1 for detailed genotypes. Scale bar = 20 µm (top and middle), 10 µm (bottom), in this and other panels. (**B**) Probability distribution of the angular position of VA1d-ORN axons in each condition at 30h APF. Population mean ± s.e.m. (**C**) Average angular position of VA1d-ORN axons in each condition. Same data as in (B). Circles indicate the averages of individual antennal lobes; bars indicate the population mean ± s.e.m. (**D**) Percentage of VA1d-ORN axons overlapping with VA1d-PN dendrites in adults. Circles indicate the average of individual antennal lobe; bars indicate the population mean ± s.e.m. (**E** to **H**) Same as (A to D), but for DA4l-ORNs and DA4l-PNs. The three manipulation conditions are: (1) *Sema-2b* RNAi; (2) *Toll-6* RNAi, *Toll-7* RNAi, and Sema-2b overexpression; (3) *Toll-6* RNAi and *Toll-7* RNAi. Note that due to limitations on the reagents, the ORN signals from the top row result from a combination of three ORN types: DA4l, DA4m, and DC1, all of which take a similar trajectory (Fig. S3). (**I–L**) Same as (A–D), but for DL4-ORNs and DL4-PNs. The two manipulation conditions are: (1) Sema-2b overexpression; (2) *Toll-6* RNAi, *Toll-7* RNAi, and Sema-2b overexpression. (**M–P**) Same as (A–D), but for DC3-ORNs and DC3-PNs. The two manipulation conditions are: (1) *Toll-7* RNAi; (2) *Toll-7* RNAi and *Sema-2b* RNAi. (**Q**) Percentage of ORN-PN mismatch in adults as a function of the absolute angular changes in ORN axon trajectory at 30h APF. The black dot indicates wild type in each ORN type, which is the origin (*x* = 0, *y* = 0) in the plot by definition. The dash line indicates the linear fit. Pearson correlation coefficient = 0.88; p = 3.6 × 10^−4^.

Take the VA1d-ORNs as an example (Fig. 5, A–D). Using three genetic manipulation strategies that combinatorially manipulated the expression levels of Sema-2b and the Toll proteins in VA1d-ORNs, we altered the ORN trajectories during development to three different distributions deviating from the wild-type distribution (Fig. 5, A top row, B and C). In adults, we observed different degrees of mismatching between the VA1d-ORNs and VA1d-PNs (Fig. 5, A bottom rows and D). We repeated this type of experiments in three other ORN types and observed different degrees of ORN-PN mismatching (Fig. 5, E–P).

When we manipulated the expression levels of Sema-2b and the Toll proteins, it is possible that we not only changed the trajectory of ORN axons but also other factors that might affect ORN-PN partner matching. The following evidence suggest that ORN-PN mismatching we observed is largely due to the change of ORN trajectories. In all cases, the portion of ORNs that mismatched with cognate PNs in adults were likely the portion of ORNs whose axons deviated from the wild-type position during development. For example, comparing ‘Manipulation #1’ to ‘wild type’ (Fig. 5A first two columns), the VA1d-ORN trajectory partly deviated counterclockwise during development (Fig. 5A top row); in adults, the VA1d-ORNs that mismatched with VA1d-PNs also moved counterclockwise (appeared as moving leftward in the lower rows). The fact that data from all the manipulation conditions followed this trend strongly suggests that synaptic partner matching is most likely due to trajectory changes, rather than to other effects caused by the change in the expression levels of Sema-2b or Tolls (which presumably happens in all ORN axons manipulated, trajectory changed or not).

Furthermore, when grouping all the data, we observed a strong positive correlation between the accuracy of ORN axon trajectories during development and the accuracy of ORN-PN matching in adults (Fig. 5Q), which means that the further ORN axons deviate from their normal positions, the more severe ORN-PN mismatch occur. These data support our working model that each ORN axon searches for synaptic partners within a narrow stripe near its axon trajectory on the antennal lobe surface, approximating a 1D space.

## Discussion

In this work, we discovered that the repeated use of the dimensionality reduction principle simplifies the synaptic partner matching problem in the assembly of the fly olfactory circuit: for each ORN type, instead of selecting 1 out of 50 PN types in a 3D volume, it only needs to select 1 out of a few PN types along a 1D trajectory (Fig. 4). A linchpin of our work has been the collection of genetic drivers that label many individual PN and ORN types across development. While single-cell-type labeling in the adult fruit fly is becoming a routine (Meissner et al., 2023; Tirian & Dickson, 2017), genetic drivers that consistently label specific cell types across development are more difficult to generate because of the dynamic nature of gene expression throughout development. Our work shows that such drivers, once generated, allow one to systematically and reliably examine the anatomical features of the same neurons at high resolution across development. This led to the discovery of PN dendrites surface extension (Fig. 1) and coincidence of partner PN dendrites and ORN axons at the surface (Fig. 2), two key bases for our model. The same drivers also allowed us to simultaneously manipulate the expression levels of multiple genes in specific cell populations across development (Fig. 3 and Fig. 5), which serves to test the model from various angles. A systematic approach of generating cell-type specific drivers throughout development (Chen et al., 2023) can propel mechanistic understandings of more developmental processes in the future.

In principle, searching for synaptic partners in a lower dimensional space reduces simultaneous choices at any given time, and thus could increase wiring accuracy and robustness. Indeed, some circuits are naturally organized following the dimensionality reduction principle, as in the case of fly optic lobe, vertebrate retina, and vertebrate olfactory bulb discussed earlier (Mori & Sakano, 2011; Sanes & Zipursky, 2020). In other circuits where synaptic targets are seemingly distributed in 3D space, the dimensionality reduction principle may nevertheless apply as in the fly antennal lobe. For example, axons of callosal projecting neurons in the mammalian cortex not only target the appropriate cortical area in the contralateral hemisphere but also terminate at specific layers (Pal et al., 2024; Wise & Jones, 1976). During development, these axons first navigate via the corpus callosum to the appropriate cortical area before ascending to specific layers (Wise & Jones, 1976; Zhou et al., 2013), converting a 3D target selection problem into sequential 2D and 1D problems. Thus, dimensionality reduction might be a widely used strategy for selecting synaptic partners in developing nervous systems.

What molecular mechanisms might be involved in executing the dimensionality reduction strategy? As is evident from the fly olfactory circuit, coordinated patterning of pre- and post-synaptic partners is required. First, PNs must target dendrites to specific 2D areas of the antennal lobe surface according to their types.

Semaphorins and leucine-rich-repeat cell-surface proteins have been shown to instruct global targeting and local segregation of PN dendrites, respectively (Komiyama et al., 2007; Hong et al., 2009; Sweeney et al., 2011). We do not know what mechanisms ensure that all PN types extend at least part of their dendrites to the antennal lobe surface, and what cause PN dendrites and ORN axons that target interior glomeruli to descent after they first contact each other at the surface. Our rerouting experiments (Fig. 3) suggest that the latter process likely involves competition with neurites from neighboring glomeruli. Second, ORN axons must choose specific trajectories (1D stripes on the antennal lobe surface) according to their types. Sema-2b has been shown to play an instructive role (Joo et al., 2013) and we further implicated Toll receptors in this study, but more molecules are likely required to fully specify ORN axon trajectories. Third, along the chosen 1D trajectory, ORN axons must select dendrites from one out of several PN types to form synaptic connections. Homophilic attraction molecules like teneurins play a role in this process (Hong et al., 2012; Xu et al., 2024) but more molecules are likely involved. We note that the dimensionality reduction strategy also enables combinatorial use of wiring molecules; for example, synaptic target selection molecules can be combined with different trajectory specification molecules so that they can be reused along spatially segregated 1D trajectories. Finally, all wiring molecules discussed above are evolutionarily conserved from invertebrates to mammals, raising the possibility that they also participate in dimensionality reduction in the wiring of the more complex mammalian brain.

## Acknowledgement

We thank the labs of Gerry Rubin, Norbert Perrimon, Yoshi Aso, Tzumin Lee and Takahiro Chihara as well as the Bloomington *Drosophila* Stock Center, the Vienna *Drosophila* Resource Center, and the KYOTO *Drosophila* Stock Center for fly stocks, Heather Dionne and Gerry Rubin for many enhancer plasmids, and members of the Luo laboratory, especially Dan Paderick and Tom Hindmarsh Sten, for helpful discussions. We also thank Junjie Luo for helpful discussion on making genetic drivers.

## Author contributions

CL, ZL, and LL conceived of the project. CL and ZL performed all of the experiments and analyzed the data. CL, ZL, and LL jointly interpreted the data and decided on new experiments. CX and KW assisted in cloning and IHC. DJL, KW, CNM, QX, TL, and HL assisted in the generation of transgenic flies. CL, ZL, and LL wrote the paper, with inputs from all other co-authors. LL supervised the work.

## Data and materials availability

All data are available in the main paper and the supplementary materials. Any additional information is available upon requests to the corresponding author.

## Supplementary Materials

**Fig. S1.**
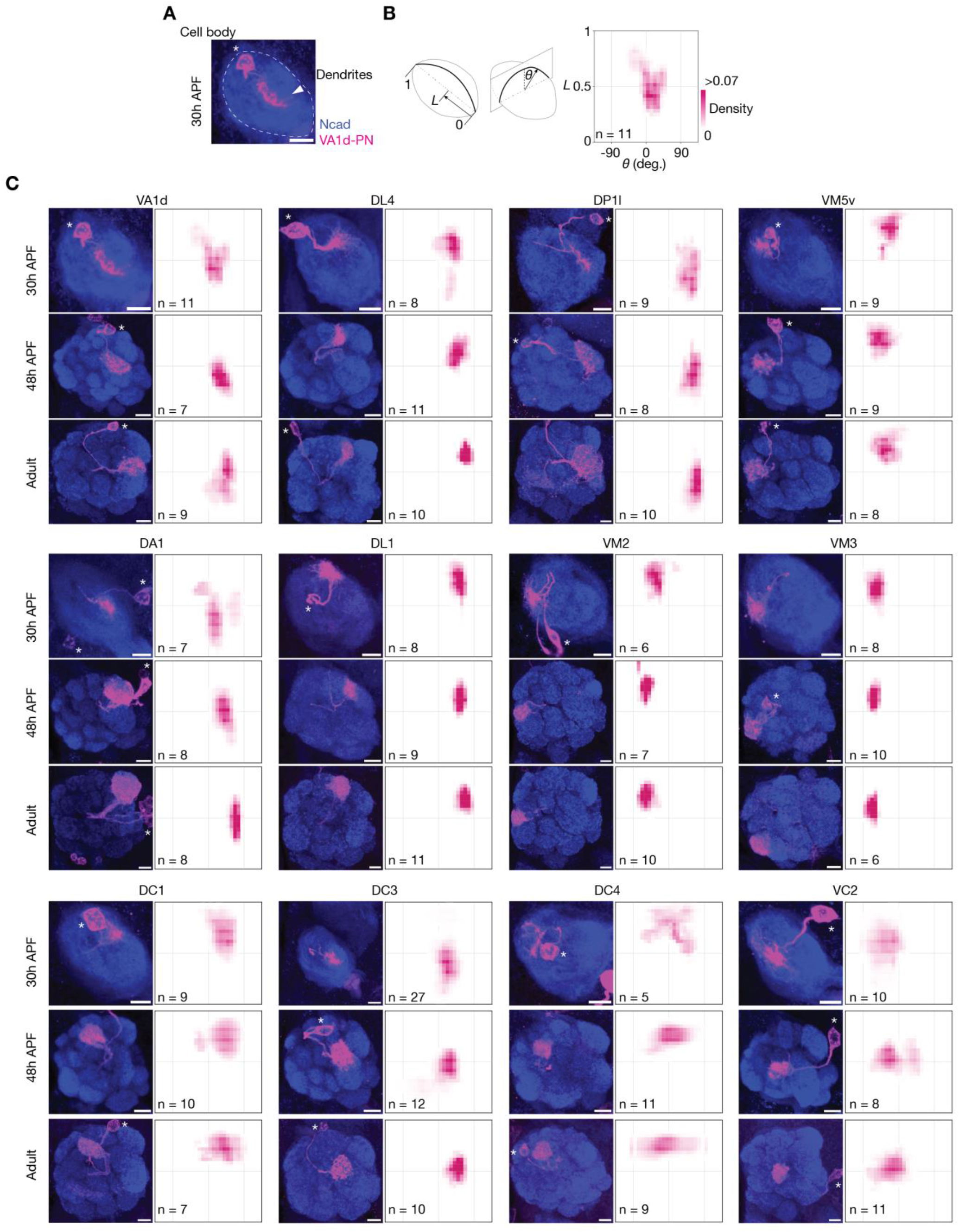
Characterization of the genetic drivers labeling specific PN types across developmental stages. (**A**) Maximum projection of horizontal optical sections of antennal lobes at 30h APF, showing dendrites of VA1d-PN (magenta, labeled by a membrane-targeted GFP driven by a genetic driver). The arrowhead indicates PN dendrites. *, PN cell body. Scale bar = 10 *µ*m in this and all other panels. (**B**) Probability distribution of VA1d-PN dendritic pixels projected onto the antennal lobe surface, averaged using data from 11 antennal lobes. The 2D antennal lobe surface is flattened and decomposed onto two axes: the *x*-axis represents the angle, *θ*, of each vertical plane and the *y*-axis represents the position, *L*, along the long axis of the antennal lobe. Schematic definition of *θ* and *L* on the left. Same heatmap scales are used for all PN types. (**C**) Same as (A) and (B), but for different genetic drivers each labeling a specific PN type across developmental stages. Note that the dendritic locations projected onto the antennal lobe surface of each PN type during development approximates their future glomerular positions in adults (Fig. 1H).

**Fig. S2.**
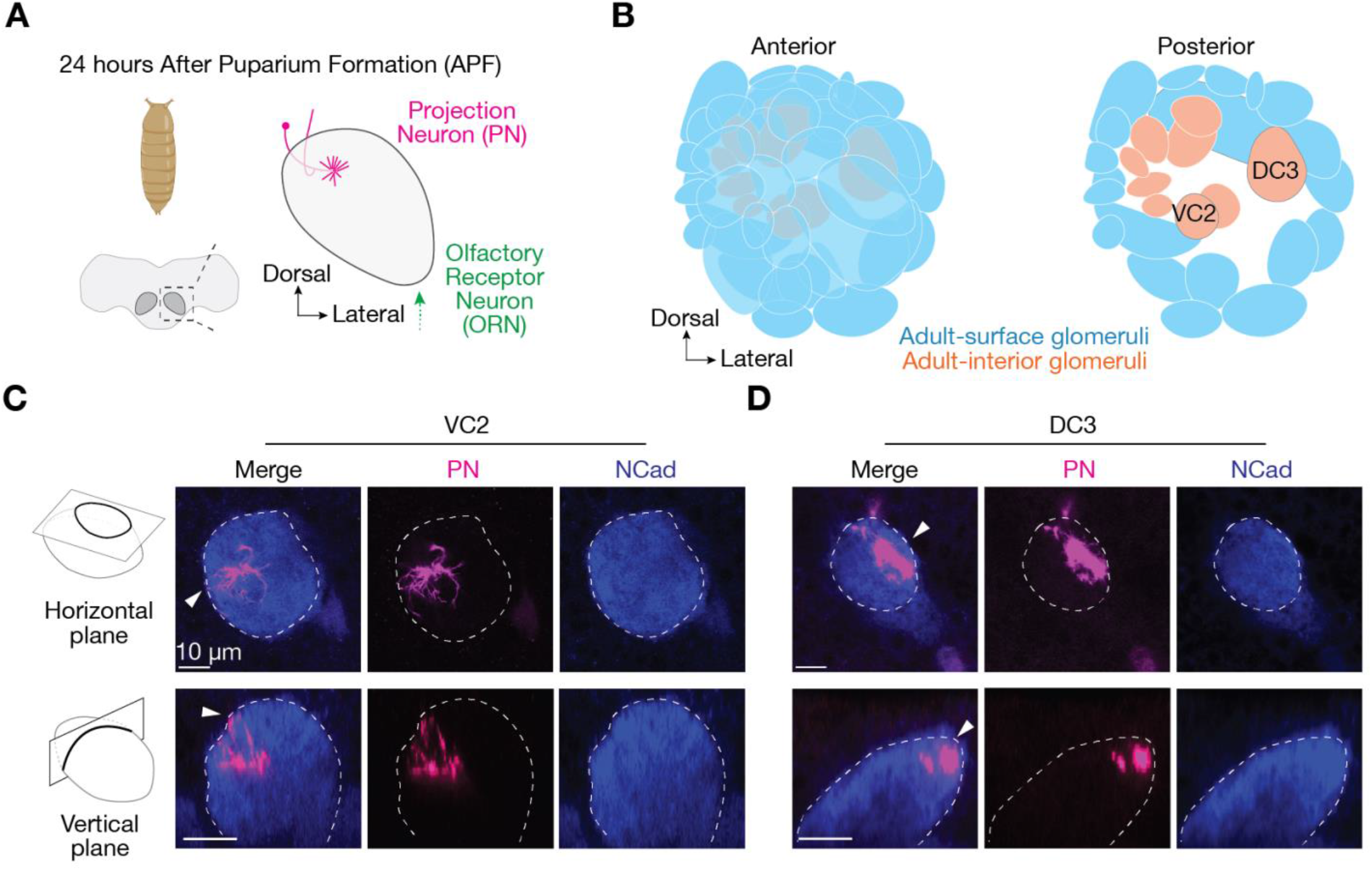
Adult-interior PN dendrites innervate the antennal lobe surface before cognate ORN axons entering the antennal lobe. (**A**) *Drosophila* brain and antennal lobe schematics, at 24h APF. Antennal lobes are highlighted in dark grey surrounded by dash squares and magnified to the right. At 24h APF, PN dendrites (magenta) innervate similar positions as in adults and most ORNs (green) have not entered the antennal lobe. For the two ORN types discussed in this figure, axons of VC2-ORNs, originating from the maxillary palp, do not reach antennal lobe until 32h APF (Sweeney et al., 2007); axons of DC3-ORNs, one of *amos*+ ORNs, do not reach antennal lobe until 28h APF (Okumura et al., 2016). (**B**) Adult antennal lobe schematic labeling the VC2 and DC3 glomeruli. (**C**) Optical sections showing dendrites of adult-interior VC2-PNs viewed from the horizontal plane (top row) and the vertical plane (bottom row) of the antennal lobe in at 24h APF. Dash lines outline the antennal lobe neuropil stained by the N-cadherin antibody (blue). Arrowheads indicate PN dendrites extending to the antennal lobe surface. (**D**) Same as (C), but for DC3-PNs. The results from (C) and (D) show that the surface extension of adult-interior PN dendrites during development is not a result of cognate ORN and PN interactions. Scale bar = 10 *µ*m.

**Fig. S3.**
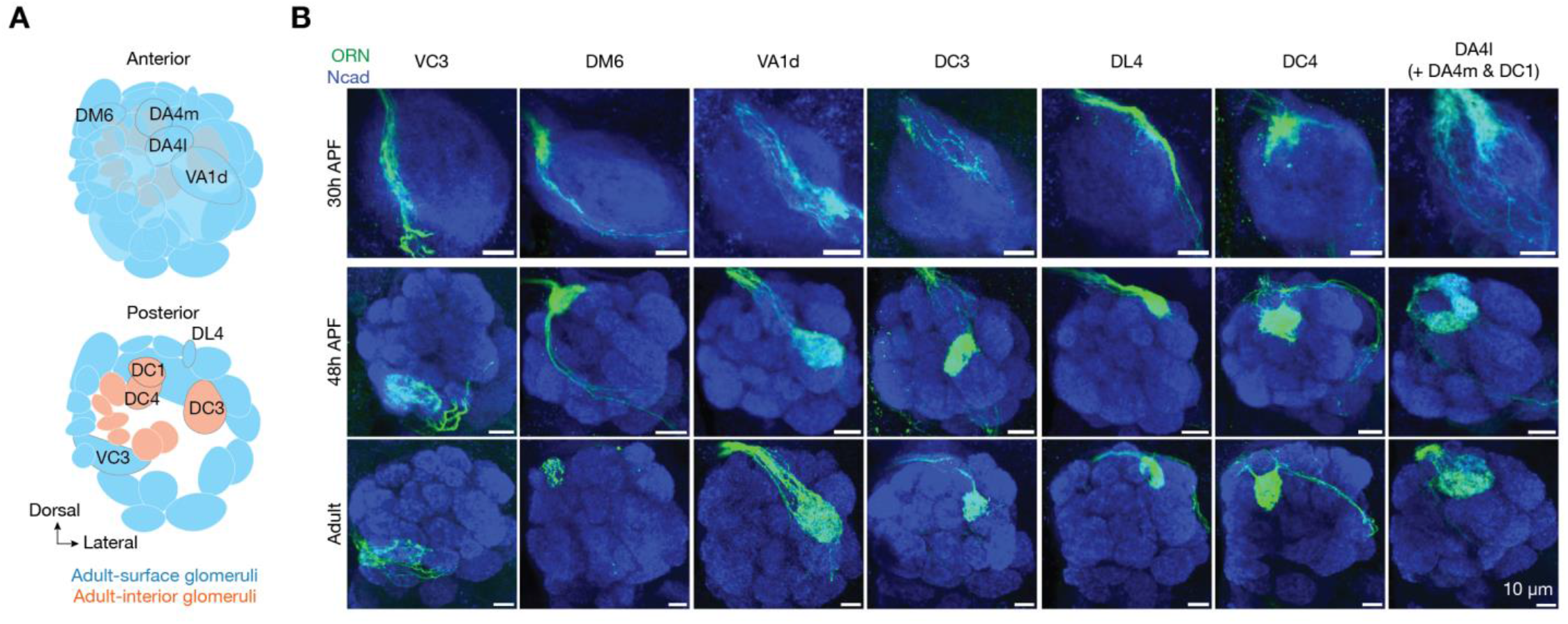
Characterization of the genetic drivers labeling specific ORN type(s) across developmental stages. (**A**) Adult antennal lobe schematic labeling nine glomeruli, corresponding to the nine ORN types shown in (B). (**B**) Genetic drivers labeling specific ORN type(s) (green, labeled by a membrane-targeted GFP) at 30h APF (top), 48h APF (middle) and in adults (bottom). Maximum projection of horizontal optical sections of antennal lobes are shown. Scale bar = 10 *µ*m. Note that all the genetic drivers most strongly label a single ORN type except for the rightmost driver, which strongly labels three ORN types.

**Fig. S4.**
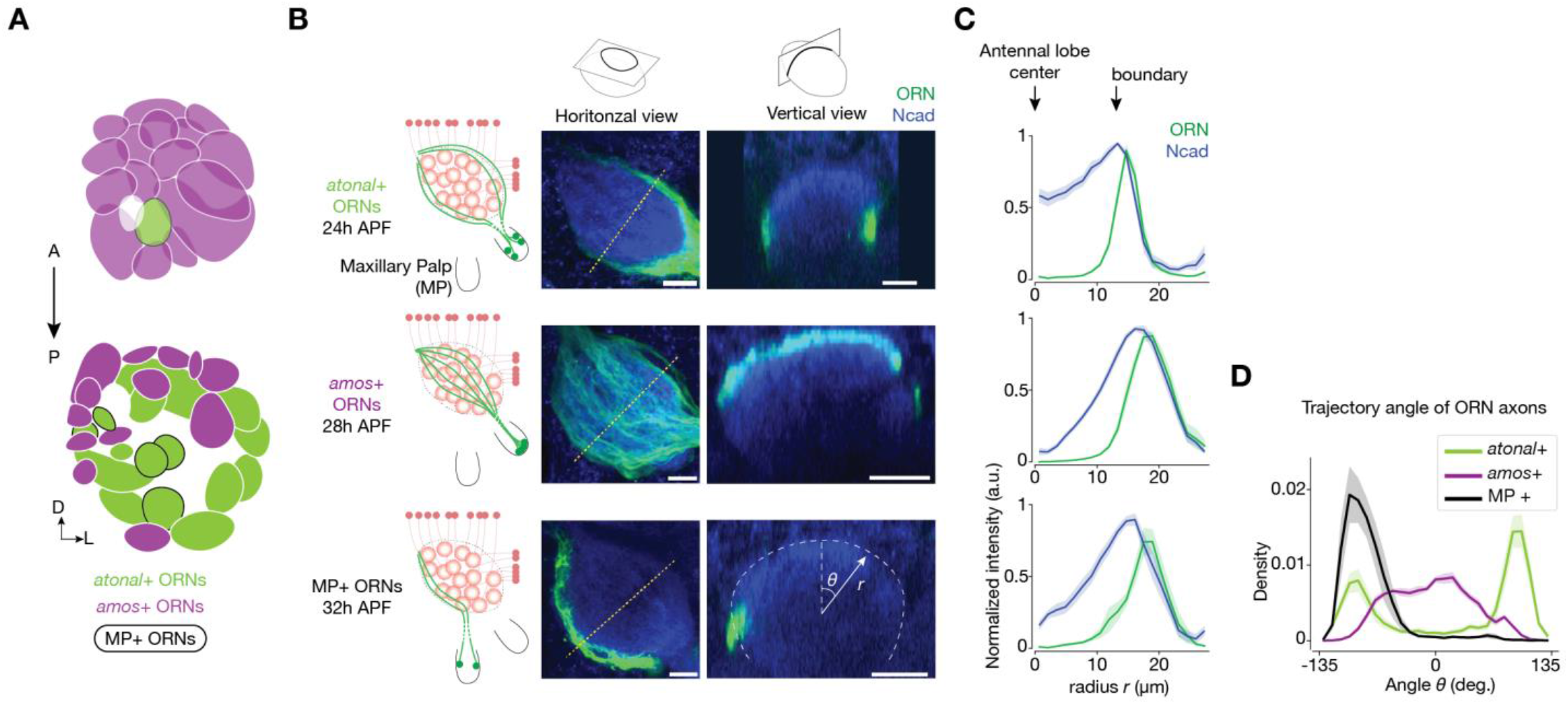
ORN axons navigate along the antennal lobe surface and collectively cover the entire anterior surface of the antennal lobe. (**A**) Adult antennal lobe schematic with different ORN groups color-coded. Purple: *amos*+ ORNs; light green: *atonal*+ ORNs; light green with black circles: *atonal*+ ORNs whose cell bodies reside in the maxillary palp. White glomeruli mean no available data. ORN axons from each group enter the antennal lobe at different time windows. (**B**) Left column, three groups of ORNs are separately labeled (green, labeled by a membrane-targeted GFP driven by separate genetic drivers) when they enter the antennal lobe. Middle column, maximum projection of horizontal optical sections of antennal lobes. Yellow dotted lines indicate the intersections with the vertical planes shown on the right. Right column, single optical sections from the vertical plane, reconstructed from 3D image volumes where optical sections were taken horizontally. Same antennal lobes for the middle and right columns. Bottom right: the dash line outlines the antennal lobe neuropil stained by the N-cadherin antibody (blue). Trajectory angle, *θ*, and radius, *r*, are defined. Scale bar = 10 *µ*m. (**C**) Signal intensities of the ORN axons and NCad in each of the three ORN groups, as a function of the antennal lobe radius. All the curves normalized to the NCad signal from each group. Only data from angular bins that have higher-than-background ORN signals were averaged. Note that in each panel, the ORN peak coincides with the right edge of the NCad peak, indicating that ORN axons navigate along the antennal lobe surface. For all groups, n ≥ 10. (**D**) Probability distribution of the angular position of ORN axons from each group. Population mean ± s.e.m. Note that the three curves combined cover angular positions ranging from –135° to +135°, which corresponds to the entire anterior surface of the antennal lobe.

**Fig. S5.**
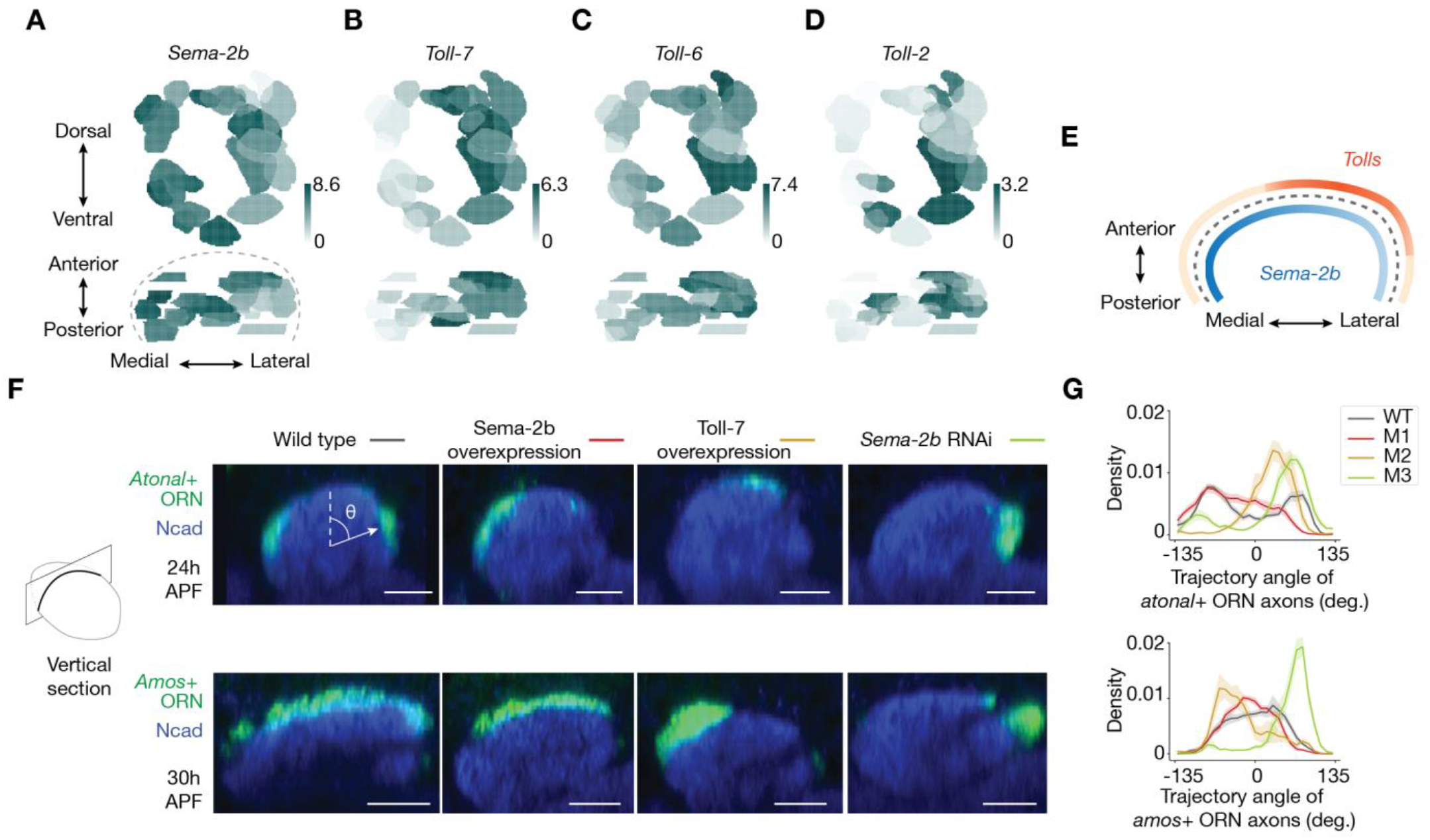
Altering the expression level of Sema-2b or the Tolls affects ORN axon trajectories. (**A**) Single-cell RNA sequencing data showing the expression level of *Sema-2b* in different ORN types at 24h APF. Heat map units: log_2_(CPM+1). CPM: counts per million reads. Top: adult glomerular map viewed horizontally. Bottom, adult glomerular map viewed vertically, along the ventral-to-dorsal axis of the antennal lobe. Note that the Sema-2b expression level is higher on the medial side than the lateral side. Data adapted from previous work (McLaughlin et al., 2021). (**B–D**) Same as (A), but for the expression levels of *Toll-7, Toll-6*, and *Toll-2*, respectively. Note that the expression level of *Tolls* is higher on the anterior-lateral side of the antennal lobe. (**E**) Schematic of the developing antennal lobe from a vertical view, with the ORN expression levels of Sema-2b (blue) and the Tolls (orange) inferred from data in (A–D). These mRNA expression levels are consistent with their corresponding proteins (Joo et al., 2013; Ward et al., 2015). This supports the working model that the angular trajectory of ORN axons is combinatorially controlled by the expression levels of Sema-2b and the Toll-family proteins in ORNs. (**F**) Single optical sections showing *atonal*+ ORN axons (top) and *amos*+ ORN axons (bottom) from a vertical view during development. Previous studies have shown that the transcription factors *atonal* and *amos* are differentially expressed in distinct groups of ORNs, with *atonal*+ ORN axons entering the antennal lobe earlier than *amos*+ ORN axons (Okumura et al., 2016). Here, we used *atonal-GAL4* and *amos-GAL4* genetic drivers to specifically label these two separate ORN groups and performed genetic manipulations within each group. Trajectory angle, *θ*, is defined in the top-left panel. Left column represents wild-type condition, other columns represent different manipulation conditions. Scale bar = 10 µm. (**G**) Probability distribution of the angular position of *atonal*+ ORN axons (top) and *amos*+ ORN axons (bottom) in each condition. Population mean ± s.e.m. For all genotypes, n ≥ 8. Note that the deviation directions of ORN axons are consistent with the map shown in (E). For example, in manipulation #1 (M1; red lines in (G)), Sema-2b is overexpressed, leading ORN axons rerouting medially where *Sema-2b* expression level is endogenously high (left side of (E)). Same is true for the other two manipulations (M2 = Toll-7 overexpression; M3 = *Sema2b* RNAi).

**Fig. S6.**
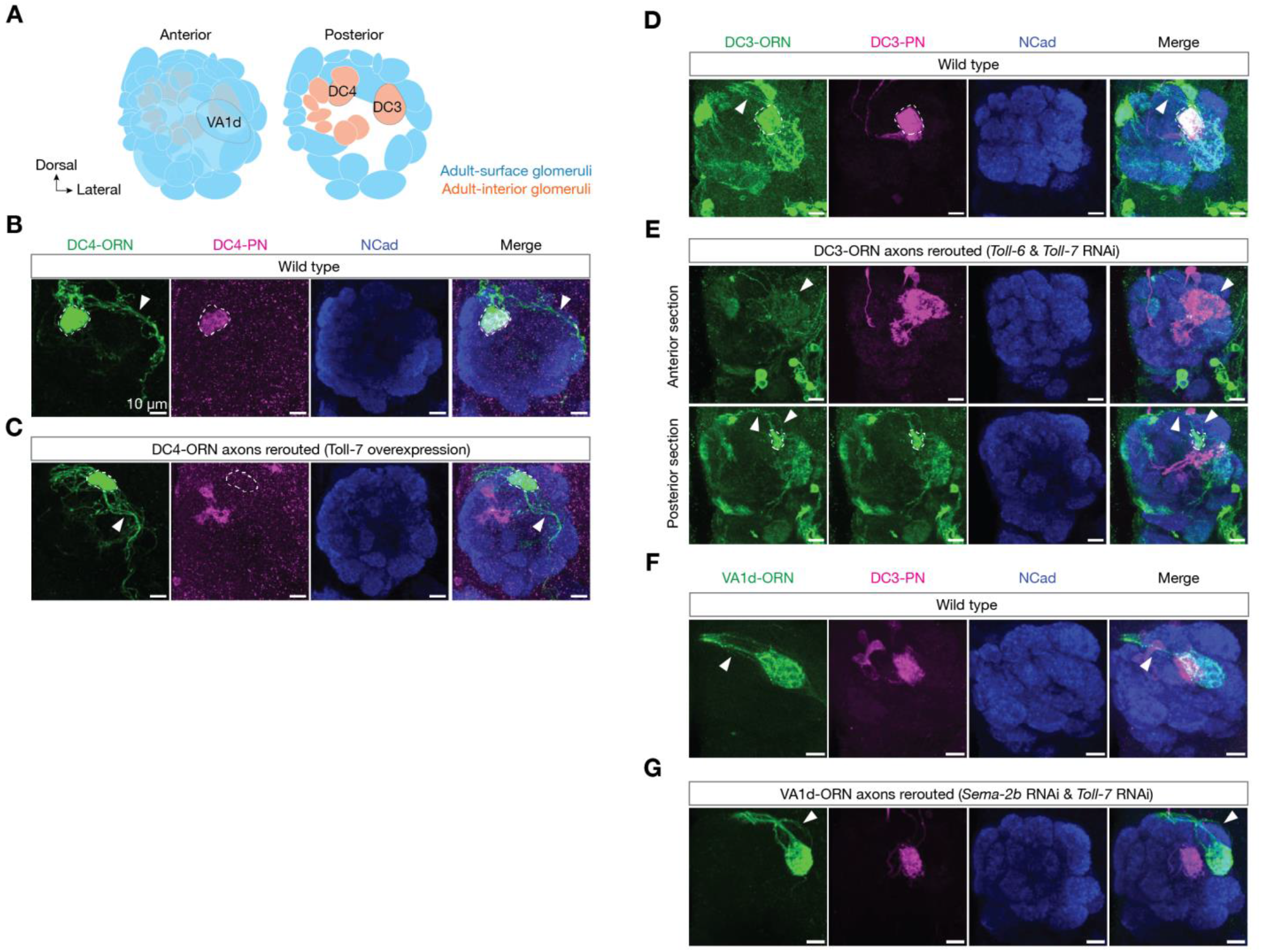
Analysis of the rerouting of DC4-ORN and DC3-ORN axons. (**A**) Adult antennal lobe schematic, highlighting three glomeruli shown in (B–F). The VA1d-glomerulus is exterior to the DC3-glomerulus. (**B**) Maximum projection of horizontal optical sections of the same antennal lobes from a wild-type brain. Dash lines outline the DC4-ORN axon terminal arborization. Arrowheads indicate DC4-ORN axons. (**C**) Same as (B), but with DC4-ORN axons rerouted by over-expressing Toll-7 in DC4-ORNs using a GAL4 driver that mainly labels DC4-ORNs in the antennal lobe (see Table S1 for the detailed genotype). Note the medial shift of DC4-ORN axons in the manipulation brain compared to the wild-type brain, leading to the complete mismatch of DC4-ORN axons and DC4-PN dendrites. (**D, E**) Same as (B) and (C), but for DC3-ORNs. (**F, G**) Same as (D) and (E), but for analyzing DC3-PNs with VA1d-ORN axon rerouted. When viewed from the vertical planes, DC3-PN dendrites are on the antennal lobe surface in conditions (E) and (G) (see Fig. 3H and 3J). Scale bar = 10 *µ*m.

### Materials and Methods

#### Fly husbandry

Flies were raised on standard cornmeal medium at 25°C with a 12-h light and 12-h dark cycle. To increase the expression level of transgenes, flies from all genetic perturbation experiments (including the control groups) were moved to 29°C shortly before puparium formation. See Table S1 for detailed genotypes of each experiment.

#### Immunohistochemistry

The procedures used for fly dissection, brain fixation and immunostaining were described previously (Xu et al., 2024). For primary antibodies, we used rat anti-DNcad (1:30, from DSHB, RRID # AB_528121), chicken anti-GFP (1:1000, from Aves Labs, RRID # AB_10000240) and rabbit anti-DsRed (1:500, from Takara Bio, RRID # AB_10013483). Janelia Fluor (JF) HaloTag dyes (JF646-HaloTag) is a gift from the L. Lavis lab (Grimm et al., 2017) and was used to stain for Halo (1:2000).

#### Molecular cloning and generation of transgenic flies

To generate QF2, GAL4DBD and LexADBD lines, we used pENTR/D-TOPO vectors with different enhancer insertions (gifts from G. Rubin lab) as entry vectors for Gateway cloning into the *pBPQF2Uw, pBPZpGAL4DBDUw*, or *pBPLexADBDzpUw* vectors using LR Clonase II Enzyme mix (Invitrogen, 11791020), respectively. *pBPQF2Uw* was made using NEBuilder HiFi DNA assembly master mix (New England Biolabs) to replace the GAL4 on *pBPGAL4*.*2Uw-2* vector (Addgene #26227) with QF2 from *pBPGUw-HACK-QF2* (Addgene #80276). *pBPZpGAL4DBDUw* is from Addgene (#26233). *pBPLexADBDzpUw* is a gift from Gerry Rubin lab. The resulting constructs were sequencing verified and inserted into *VK00027, attP2* or *attP86Fb* landing sites by Bestgene. *Slp1-T2A-AD* and *Arc1-T2A-AD* were generated using CRISPR mediated knock-in as previously described (Xie et al., 2021). In brief, genomic sequences flanking the targeted insertion sites were amplified and inserted into the *T2A-p65AD* vector to generate donor vectors *TOPO-slp1-T2A-p65AD-P3-RFP* and *TOPO-arc1-T2A-p65AD-P3-RFP*. gRNA target sequences were selected by the flyCRISPR Target Finder tool and were cloned into *pU6-BbsI-chiRNA* (Addgene #45946) to make gRNA vectors. The donor vector and the gRNA vector were co-injected into *vas-Cas9* embryos. *Mz19-AD*^*G4HACK*^, *Pebble-AD*^*G4HACK*^, and *AM29-DBD*^*G4HACK*^ was generated by injecting *pBPGUw-HACK-G4-split-p65AD or HACK-G4-split-DBD* (Xie et al., 2019) into *Mz19-GAL4, Pebble-GAL4*, and *AM29-GAL4* embryos (with *Cas9*), respectively.

#### Imaging

Immunostained brains were imaged using a laser-scanning confocal microscope (Zeiss LSM 780). Images of antennal lobes were taken as confocal stacks with 1-mm-thick sections. Representative single sections were shown to illustrate the arborization features of ORN axons and PN dendrites, with brightness adjustment, contrast adjustment, and image cropping done in ImageJ.

#### Reconstructing vertical image planes from 3D image volumes

To view the ORN axons and PN dendrites from a vertical perspective orthogonal to navigating ORN axons, we computationally reconstructed vertical image planes from 3D image volumes where optical sections were taken horizontally. Since ORN axons enter the antennal lobe at its ventrolateral corner and cross the midline near the dorsomedial corner of the antennal lobe, in each antennal lobe image, we fit the antennal lobe signal (NCad channel) to an ellipse and used the long axis of this fitted ellipse as an estimate of the average navigating direction of ORN axons. Each ellipse is fitted using the NCad data from the single horizontal plane with the largest NCad area.

#### Calculating the distance from PN dendritic pixels to the antennal lobe surface

The boundary of the antennal lobe in each *z*-plane was manually outlined by an experimenter based only on the NCad signal (i.e., blind to PN signals). Then the distance from every PN dendritic pixel to the antennal lobe surface was calculated by finding the shortest distance from the PN pixel to the antennal lobe surface from a 3D space, i.e., by finding the minimum value from the PN pixel to all the digital points belonging to the antennal lobe boundary throughout all the z-planes. Each PN dendritic pixel was defined by first smoothening the image using ‘gaussian blur’ (radius = 2 pixels) and then thresholding the image based on the algorithm ‘Otsu’ in Fiji. We found that this algorithm could efficiently separate the neurons of interest from the background. Irrelevant signals (such as the PN axons, cell bodies, or autofluorescence) that still persist after the above operations were manually masked out in the analysis.

#### Calculating the angular position of ORN axons

Since the angular position was defined in vertical planes, all angular positions were calculated using imaging data reconstructed into vertical planes (see above) and were averaged from vertical planes near the center of the antennal lobe long axis where measurement variance is minimum. For each of the vertical planes calculated, the centroid is the intersection point from the antennal lobe long axis. Then in each angular bin, maximum fluorescence intensities (such as from the channel of ORN axons or PN dendrites) were used after subtracting the baseline from background.

#### Calculating the percentage of ORN axons matching with PN dendrites

Pixels of ORN axons were defined using similar methods as described above (in the section of ‘Calculating the distance from PN dendritic pixels to the antennal lobe surface’). The portion of ORN axons were considered as matching with PN dendrites if they have overlapping pixels on a single *z*-plane in the image.

